# A Mouse Model with Complete Penetrance for Atrioventricular Septal Defect/Complete AV Canal

**DOI:** 10.1101/2022.02.03.478994

**Authors:** Yicong Li, Peter Andersen, Xihe Liu, Anna J. Moyer, Chulan Kwon, Roger H. Reeves

## Abstract

Complete Atrioventricular Septal Defect/complete AV canal (AVSD/ cAVC) occurs via defective formation of the Dorsal Mesenchymal Protrusion (DMP) which grows from the interior dorsal wall of the fetal heart. The DMP is derived from cells of the second heart field (SHF). AVSD occurs in about 1/10,000 births in the general population, but is 2000 times more prevalent in individuals with Down syndrome (DS). The low penetrance of AVSD in the general population remains a fundamental challenge to investigation of the genetic and developmental basis for this condition. Analysis of the etiology of this condition in mouse models is limited by the fact that the developmental events producing the DMP begin around embryonic day 9 (E9) but AVSD cannot be diagnosed until E14, and no model has been described previously in which AVSD occurs in 100% of embryos. We describe a trisomic mouse DS model with conditional mutations in the SHH pathway that produces AVSD with 100% penetrance. This new mouse model can serve as an important tool to understand the mechanisms underlying AVSD pathogenesis.

## Introduction

Congenital heart defect (CHD) has been identified as the most common birth defect; nearly 1 out of 100 newborns has a structural heart problem ^1^. Risk of CHD is further elevated to more than 40% among those with trisomy 21, the cause of DS ^2,3^. Complete Atrioventricular Septal Defect (AVSD) occurs in 20% of babies with DS, a 2000-fold increase over the general population ^4^. Patients with AVSD typically require cardiac surgery during their first year of life. They remain at high risk of arrhythmias, endocarditis, stroke, congestive heart failure, and other conditions ^3^. Despite the high morbidity and mortality of AVSD and the greatly increased incidence in people with DS, little is known about the trisomic genes whose over-expression contributes to these increased risks.

More than 20 mouse models of trisomy 21 have been created by duplication of segments of mouse chromosomes 16 (Mmu16), Mmu10, and Mmu17 that are orthologous to human chromosome 21 (Hsa21) ^5^. In addition, humanized DS mouse models containing portions of Hsa21 as a freely segregating chromosome have been created to assess various DS-related phenotypes ^6,7^. While CHD occurs in a number of these models, none of them presents with complete AVSD nor do they occur with complete penetrance in the offspring (100% affected). The Dorsal Mesenchymal Protrusion (DMP) is responsible for combining the cap of the primary atrial septum and superior AV cushion during heart development ^8,9^. The most severe form of AVSD, sometimes referred to as complete AV canal (cAVC), occurs due to a failure to form the DMP. In contrast to the greatly increased risk of cAVC in DS, this condition has never been shown to occur in trisomic mouse models of DS. Unfortunately, the term AVSD is used to describe several different conditions resulting in communication between heart chambers that differ structurally in different developmental fields, demonstrating differences in etiology. The non-specific use of this term has led to considerable confusion about the origins of this structural defect. *Herein, we use AVSD to refer to a complete AV canal due to failure of DMP formation as occurs in DS*.

Cardiac progenitor cells are specified during gastrulation and form two heart fields, referred to as the first heart field (FHF) and the second heart field (SHF). The FHF gives rise to the right ventricle and part of atria, while the SHF contributes to the outflow tract, right ventricle, and part of atria ^10^. Several lineage tracing studies showed that the DMP is also derived from the SHF, and its development is regulated by Sonic Hedgehog (SHH) signaling ^9,11-14^. Mouse models in which SHH growth factor signaling is disrupted in the SHF show AVSD with incomplete penetrance (i.e., not all embryos are affected). Previous studies have demonstrated an attenuated response to SHH signaling in trisomic mouse DS models, documented in multiple trisomic tissues ^15,16^. Thus, alterations in SHH signaling may contribute to multiple features of DS, including AVSD. To date, however, none of the available models is completely penetrant, and the presence or not of AVSD can only be determined by histology at E14.5. Since critical steps in the process forming the DMP occur around E9.5, a mouse model in which it is known absolutely which embryos will develop AVSD is an important tool for understanding the etiology of AVSD.

## Results

### A mouse model with completely penetrant AVSD (cAVC)

The critical linkage between SHH signaling and heart development, the increased frequency of heart defects in DS mouse models and of AVSD particularly in babies with DS, plus the attenuated SHH response in DS suggested an approach to the development of a highly penetrant AVSD model. A somewhat involved breeding strategy was necessary to introduce the necessary genetic modifications. The *Isl1* Homeobox transcription factor is widely expressed in SHF cells and *Smo* is the canonical SHH pathway regulator. We reduced hedgehog signaling in SHF progenitor cells by expression of Isl1-Cre in mice that were heterozygous or homozygous for a floxed allele of *Smo* ^17^. Cre is “knocked in” to the *Isl1* gene, creating a null allele that is homozygous lethal, and therefore is carried as a heterozygous allele (Isl1^*Cre/+*^)^18^.

Dp(16Lipi-Zbtb21)1Yey mice (Dp16) carry a direct duplication of the distal portion of mouse chromosome 16 (MMU16) that is conserved with HSA21^19^. We crossed Isl1^*Cre/+*^ to Dp16 mice that had been made homozygous for the *Smo*^*fl*^ allele (which express Smo normally in the absence of Cre) to produce trisomic and euploid backgrounds. Approximately 1/10 embryos recovered at E14.5 carried an *Isl1*^*Cre*^ allele, were homozygous for the desired conditional *Smo* deletion and were recovered on the trisomic background. All mice were homozygous for a TdTomato reporter that was activated by Isl1-Cre - mediated excision of a lox-stop-lox cassette.

Five genotypes of mice were assessed, all segregating Isl1-Cre and a lox-stop-lox TdTomato marker gene (TdT reporter): euploid *Isl1*^*Cre/+*^ controls; trisomic Dp16 mice; trisomic mice heterozygous for a floxed allele of *Smo* (*Smo*^*fll+*^); euploid mice homozygous for a floxed allele of *Smo* (*Smo*^*fl*/*fl*^); and Dp16;*Smo*^*fl*/*fl*^ (“Triple” mice) (**Table 1**).

**Table 1.** Congenital heart defect frequency at E14.5 by genotype.

| Genotype<sup>a</sup> | ASD | VSD | AVSD<sup>c</sup> | Total | % CHD | % AVSD |
| --- | --- | --- | --- | --- | --- | --- |
| *Isl1*-Cre | 1 | 2 | 0 | 26 | 12% | 0 |
| *Isl1*-Cre;Dp16 | 0 | 1 | 0 | 11 | 9% | 0 |
| *Isl1*-Cre;Dp16;*Smo*<sup>fl/+</sup> | 1 | 3 | 10 | 15 | 93% | 67% |
| *Isl1*-Cre; *Smo*<sup>fl/fl</sup> | 1 | 3 | 2 | 8 | 75% | 25% |
| *Isl1*-Cre;Dp16;*Smo*<sup>fl/fl</sup> | 0 | 0 | 12 | 12 | 100% | 100% |

<sup>a</sup> All mice were carried on a C57BL/6J background for at least 6 generations. All mice carrying *Isl1*-Cre also carried a lox-stop-lox TdTomato marker transgene to detect *Isl1* positive cells based on *Isl1*-Cre expression.
<sup>b</sup> Analyzed previously at P0<sup>22</sup>.
<sup>c</sup> Complete AV canal type due to absence of DMP.

Cardiac septation is completed by E13.5, therefore we recovered embryos at E14.5. Hearts were assessed histologically in paraffin-embedded sections of embryos (**Fig. 1** and **Table 1**). Genotypes were assessed by PCR and only associated with heart condition after assessment (i.e., those scoring heart defects were blind to genotype). As in previous reports of Dp16Yey1 and the effectively equivalent Dp16Tyb1 trisomic mouse strains ^20,21^, we observed ASD and VSD but no AVSD (i.e., no cAVC; Dp16Tyb1 is reported to have AVSD, but it is not cAVC due to failure to form a DMP, i.e., not the prominent form of AVSD that occurs in DS) (**Table 1**). Septal defect frequencies in *Isl1*^*Cre/+*^ and *Isl1*^*Cre/+*^; Dp16, the euploid and trisomic controls, were consistent with previous results. Homozygous deletion of *Smo* in euploid mice produced septal defects in 75% of embryos, but only two of these (25%) were AVSD. Trisomy plus one normal and one floxed allele of *Smo* (Dp16;*Smo*^*fll+*^) produced 67% AVSD, while 100% of Dp16;*Smo*^*fl*/*fl*^ mice had AVSD, a significant difference (p<0.05). Thus we can predict with high confidence that any embryo with the *Isl1*^*Cre/+*^;Dp16;*Smo*^fl/fl^ genotype will develop AVSD well before that condition can be determined by phenotyping (i.e., histology or ultrasound).

**Figure 1.**
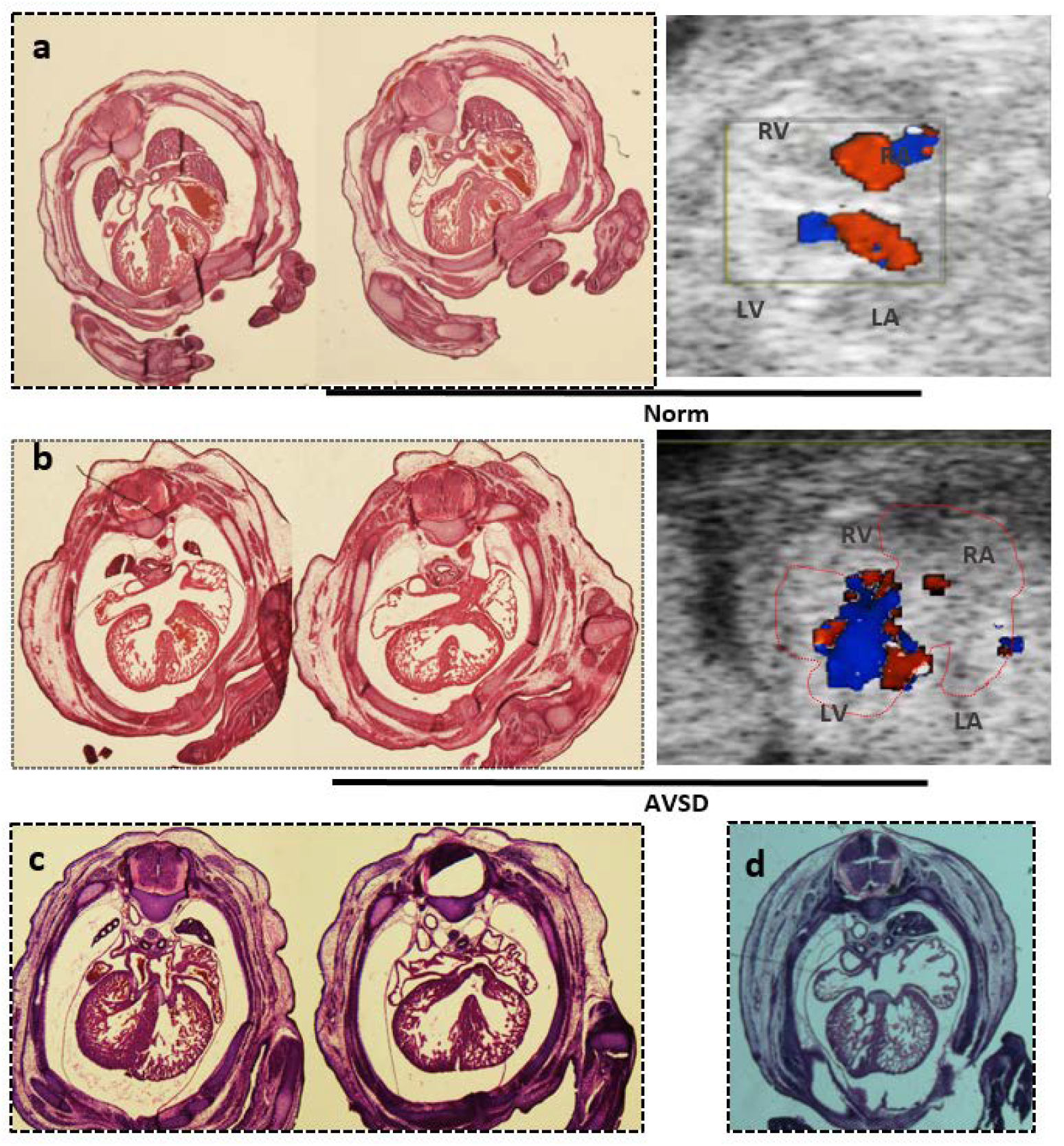
Heart defects in mouse models. Hearts were assessed by histology to identify a) normal, b) AVSD, c) VSD and d) ASD. Ultrasound shows normal blood flow in a) and mixing of pulmonary and systemic circulation in b).

## Discussion

Important events in the process of heart development can be studied in cell culture, especially using differentiating iPSCs from different genetic backgrounds, but ultimately studies of structural defects such as AVSD require an animal model. We generated trisomic mice with further reduced SHH signaling due to conditional deletion of *Smo* and found that 100% of progeny developed AVSD of the complete AV canal type (“AVSD mice”). The ability to predict which embryos are experiencing maldevelopment of the DMP at any stage of development provides the necessary model for detailed investigation.

Here we targeted early stages of DMP formation by isolating *Isl1*-expressing SHF cells migrating from pharyngeal arches that are known to serve as a microenvironment for undifferentiated SHF cells ^23^. This is based on our assumptions that (1) DMP development initiates prior to the migration and differentiation of SHF cells in the arches and (2) aSHF cells are a major cell source used to form the DMP. The former is supported by several in vivo studies demonstrating the importance of hedgehog signaling in SHF cell proliferation and DMP formation ^11,12,14,24,25^. Yet, it remains to be determined when and where DMP progenitors are specified from the pool of undifferentiated SHF cells. The latter assumption is based on several lineage tracing experiments showing that aSHF cells give rise to the DMP ^12,26^. pSHF cells, which migrate through the inflow tract of the developing heart, have also been shown to play an important role in DMP formation ^27,28^.

The essential elements for production of embryos with AVSD in this model are trisomy, introduced from a heterozygous parent, and a gene that reduces expression of the SHH pathway. We targeted *Isl1*-expressing cells using a floxed allele of *Smo* that was homozygous in one parent without Cre and heterozygous in the other to produce the desired genotype. Additional Cre regulation, e.g. with an ER-Cre, could refine questions regarding critical windows for timing of Smo expression, while Cre promoters expressed in different subsets of migrating SHF cells would support interrogation of spatial regulation by additional genes with roles in subsets of SHF cells. In any regard, the ability to study heart development in a model that will reliably develop AVSD should prove valuable to further understanding of this serious condition.

## Materials and Methods

### Animal husbandry and genotyping

Mice used in this study were maintained in an AAALAS-certified clean facility with food and water ad libitum. All strains were maintained on the C57Bl/6J background (B6J) for more than 6 generations. All procedures were approved by the Institutional Animal Care and Use Committee.

Genomic DNA extracted from tail tips and was used for genotyping. For Dp16, PCR typing was performed as described ^29^. Additional PCR typing was performed using the following primers:

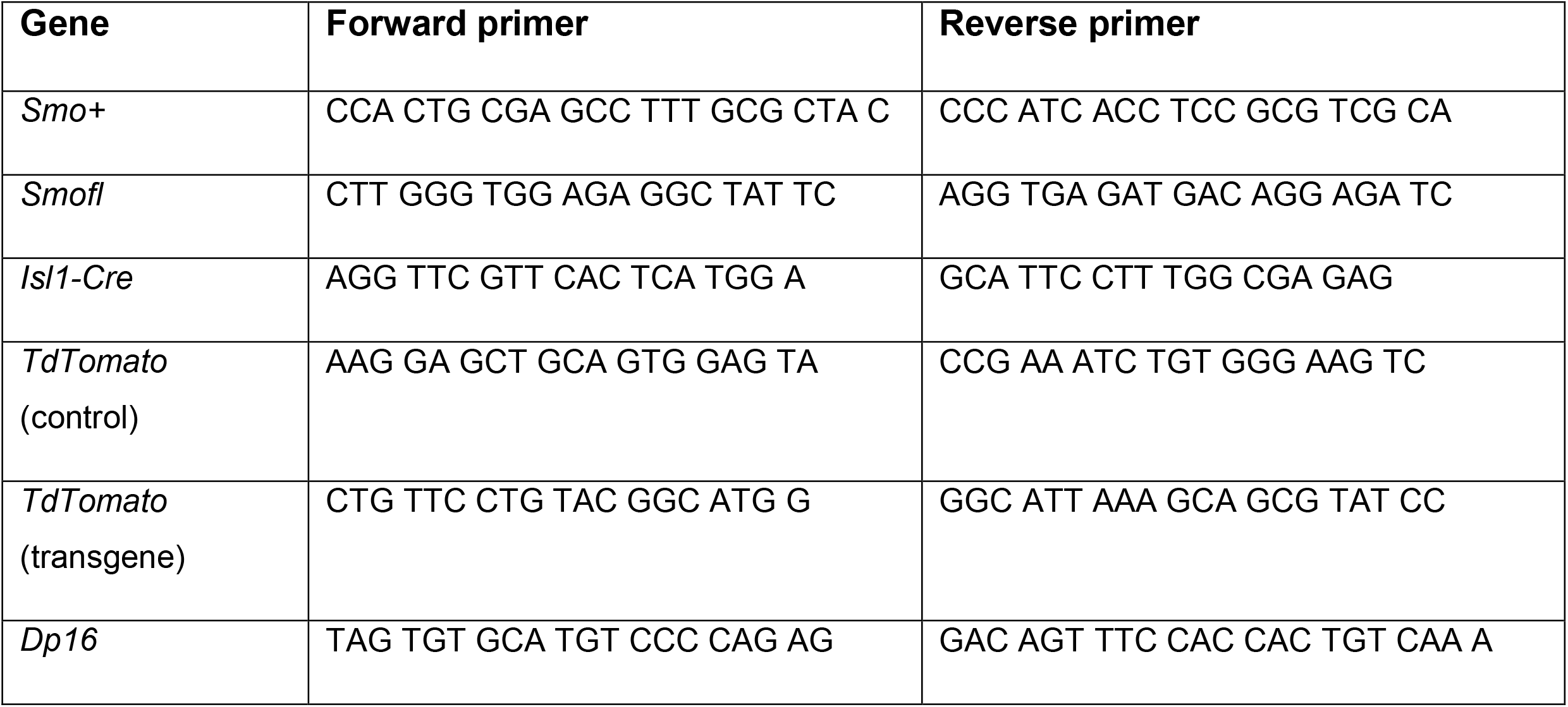

### Histology and ultrasound analysis

Embryos were collected at E14.5 post-copulatory plug, thorax was removed, fixed in 4% paraformaldehyde for two hours, dehydrated and embedded in paraffin. Fifteen um sections were taken through the block and stained with haemotoxylin and eosin for assessment.

A Vevo 2100 Echocardiography system was used for ultrasound. The uterus was exposed in E14.5 timed pregnant mothers for scanning including four chamber view, m-mode, color Doppler and blood flow direction.

### Statistical analysis

Data were processed using Microsoft Excel and GraphPad Prism 9. All values are presented as the mean ± S.E.M. Comparisons within two groups using Microsofr Excel were determined using two-tailed, unpaired t test. Analysis was performed using one-way ANOVA (nonparametric) test and followed by multiple comparisons testing which compared the mean of each group with the mean of the control group. P values less than 0.05 were considered to be significant.

## Acknowledgements

This work was supported by Public Health Service grants 1R21HD098540-01 and MPI R01HL124836 (RHR), 1F31HD098826 (AJM) and R01HD086026) (CK) and by grants from the Saving Tiny Hearts Society (CK).

